# FluNexus: a versatile web platform for antigenic prediction and visualization of influenza A viruses

**DOI:** 10.64898/2026.01.29.702696

**Authors:** Xingyi Li, Chunyan Zhou, Han Wu, Kexin Xiao, Jun Hao, Dongmin Zhao, Junnan Zhu, Yue Li, Jiajie Peng, Jia Gu, Guohua Deng, Weigang Cai, Min Li, Yan Liu, Xuequn Shang, Hualan Chen, Huihui Kong

## Abstract

Influenza A viruses continuously undergo antigenic evolution to escape host immunity induced by previous infections or vaccinations, consequently causing seasonal epidemics and occasional pandemics. Antigenic prediction and visualization of influenza A viruses are crucial for precise vaccine strain selection and robust pandemic preparedness. However, a user-friendly online platform for these capabilities remains notably absent, despite widespread demand. Here, we present FluNexus (https://flunexus.com), the first-of-its-kind, one-stop-shop web platform designed to facilitate the prediction and visualization of the antigenic change in emerging variants. FluNexus features a data preprocessing module for hemagglutinin subunit 1 (HA1) and hemagglutination inhibition (HI) data across three major public health threat subtypes (H1, H3 and H5). Meanwhile, FluNexus provides an interactive interface for online antigenic prediction and offers practical guidance for researchers. Most notably, FluNexus offers the visualization of influenza A virus antigenic evolution, providing intuitive insights into its antigenic dynamics. Specially, FluNexus proposes a novel manifold-based method for positioning antigens and antisera, generating accurate antigenic cartographies even with sparse HI data. By alleviating the programming burden on biologists, FluNexus supports more informed decision-making in vaccine strain selection and strengthens surveillance and pandemic preparedness.

**Highlights:**

- FluNexus features a data preprocessing module for HA1 and HI data spanning the H1, H3, and H5 subtypes.
- FluNexus facilitates online antigenic prediction utilizing ten state-of-the-art antigenic prediction tools, and offers practical guidance based on a comparative evaluation of their performance.
- FluNexus provides a visualization module for mapping antigenic evolution of influenza A viruses, incorporating a novel manifold-based method for antigenic cartography.

## 1 INTRODUCTION

Influenza A viruses pose a major global health threat due to their high mutation rate, broad host range, and potential to cause both epidemics and pandemics [1]. To date, 19 hemagglutinin (HA) and 11 neuraminidase (NA) subtypes have been identified [2]. Human seasonal influenza is primarily driven by H1 and H3 subtypes, which account for an estimated 3-5 million severe cases and 290,000-650,000 respiratory-related deaths annually worldwide. Beyond seasonal outbreaks, avian influenza viruses also present a pandemic threat, particularly the H5 subtype, whose widespread circulation in birds caused serious damage to poultry production, dairy industry and public human health, resulting in the culling of more than 389 million poultry, infections in over 1,000 dairy cattle herds, and more than 1,080 human cases [3–5]. Vaccination is a cornerstone strategy for controlling both seasonal and avian influenza.

However, the protective efficacy of vaccines is commonly undermined by rapid antigenic evolution, driven predominantly by mutations in the hemagglutinin subunit 1 (HA1) proteins that confer immune escape. When antigenically matched vaccines are deployed in a timely manner, they can effectively control or even eliminate circulating viruses. For example, H7N9 avian influenza viruses that caused repeated human infections between 2013 and 2017 are eliminated following the introduction of an antigenically matched H7 vaccine strain in poultry [6]. In practice, delays in updating vaccines to match antigenic changes increase the risk of outbreaks, underscoring the urgent need for rapid, high-throughput, and accurate methods to assess the antigenicity.

Conventionally, influenza virus antigenicity is assessed using serological assays such as hemagglutination inhibition (HI), which are labor-intensive, time-consuming, and low-throughput. Unlike HI assays for H1 and H3 viruses, which can be performed in biosafety level 2 (BSL-2) facilities, HI assays for H5 viruses must be conducted in BSL-3 laboratories, further **slowing** the process of data generation. These limitations hinder the capacity of traditional methods to keep pace with the growing number and antigenic diversity of influenza isolates. Advances in sequencing technologies and the increasing availability of viral genomic data facilitate the development of computational methods for fast antigenic prediction, thereby improving vaccine efficacy and strengthening pandemic preparedness. Smith et al. [7] depict the antigenic evolution of A/H3N2 using modified metric multidimensional scaling (MDS) to reveal cluster-wise, low-dimensional shifts closely aligned with genetic changes, as named as Racmacs. This methodology has been extensively refined in follow-up studies [8, 9]. Subsequent efforts have focused on developing machine learning [10–13] and deep learning-based methods [14–19] for antigenic prediction by combining HI data with large-scale sequencing data.

Despite the availability of computational methods for exploring antigenic evolution in recent years, key challenges remain. Specifically, user-friendly tools for online data preprocessing and antigenic prediction are still lacking. Moreover, the visualization of antigenic distances among variants is crucial for interpreting HI titers and assessing the accuracy of prediction results. Effective antigenic cartography empowers virologists and immunologists to rapidly understand the antigenic relationships between variants. However, most existing tools require programming expertise, which hinders their accessibility for many bench scientists without computational backgrounds. There is an urgent need for a versatile, user-friendly online platform to facilitate the systematic monitoring of antigenic evolution in emerging variants, thereby promoting more timely and evidence-based vaccine strain selection.

To address these limitations, we propose FluNexus, a first-of-its-kind web platform for antigenic prediction and visualization of influenza A viruses with a user-friendly interface. It first features interactive data preprocessing modules for HI and HA1 data. Next, FluNexus supports ten state-of-the-art sequence-based computational algorithms for online antigenic prediction, and presents a comprehensive benchmarking framework to evaluate their performance for H1, H3, and H5 subtypes in terms of discriminating antigenic variants from non-variants, predicting antigenic distances of virus pairs, and computational time complexity, thereby offering practical recommendations for users conducting influenza antigenic prediction. Moreover, FluNexus provides the visualization of antigenic map and antigenic cluster of influenza A viruses, and proposes an optimized strategy for antigenic cartography.

## 2 RESULTS

### 2.1 mPipeline and web platform overview

As a pioneer web platform for antigenic prediction and visualization of influenza A viruses, FluNexus integrates three core components-data preprocessing, online computational tools and evaluation, and antigenic visualization-into a unified, accessible framework. The architecture and implementation of FluNexus are shown in Supplementary S1, and users can navigate easily among all functional modules through the web interface. A detailed user manual is available online at https://flunexus.com/tutorial.html, providing comprehensive guidance on data input, method usage, and result interpretation. The overall workflow is illustrated in Figure 1.

**FIGURE 1.**
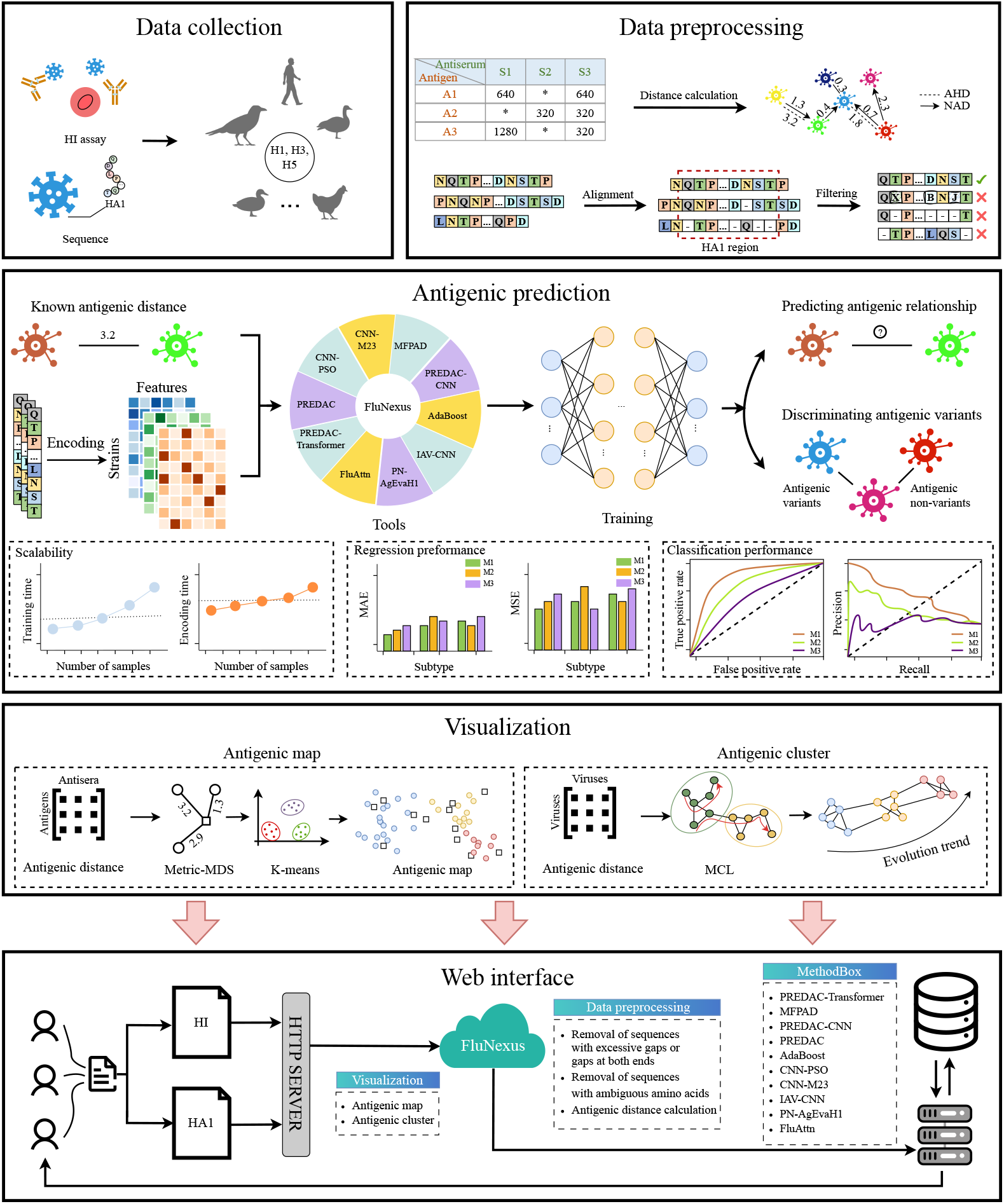
Overview of FluNexus. FluNexus comprises online data preprocessing and antigenic prediction modules. Furthermore, the platform facilitates the visualization of antigenic maps and clusters, as well as the proposed optimized cartography strategy. Ultimately, FluNexus empowers researchers with a versatile web platform for effective influenza surveillance and analysis.

### 2.2 Data preprocessing and antigenic prediction tools

To facilitate convenient access to unified HA1 and HI data for antigenic prediction or visualization, FluNexus features a data preprocessing pipline for users to process their own uploaded data on the ‘Data Preprocess’ page. For HA1 sequences, FluNexus provides three quality control options: (i) remove uncertain amino acids (e.g., B, Z, J, X), (ii) remove sequences with gaps at both ends, and (iii) remove sequences with high gap ratios. For HI data, the platform facilitates the conversion of HI titers into antigenic distances. Specifically, antigenic distances of virus pairs can be calculated using either the Normalized Antigenic Distance (NAD) [20] or the log2-transformed Archetti–Horsfall Distance (AHD) [21, 22] (METHODS). Based on the calculated antigenic distance, each virus pair can be classified as antigenically similar (distance *<* 2) or distinct (distance *≥* 2), consistent with criteria widely used in Centers for Disease Control and Prevention (CDC) or World Health Organization (WHO) surveillance reports (https://www.cdc.gov/fluview/surveillance/2025-week-52.html) and influenza antigenic cartography studies [7, 17, 23, 24].

Accurate sequence-based predictive methods are pivotal for enabling large-scale antigenic surveillance of circulating viral isolates while reducing the reliance on labor-intensive experimental assays. To this end, FluNexus provides both online and open-source implementations (https://github.com/xingyili/FluNexus-methodbox) of ten sequence-based tools for predicting antigenic distance and discriminating antigenic variants from non-variants. The integrated tools comprise PREDAC [10], CNN-PSO [14], CNN-M23 [15], MFPAD [12], PREDAC-CNN [17], AdaBoost [11], IAV-CNN [16], PN-AgEvaH1 [13], PREDAC-Transformer [18], and Flu-Attn [19] (METHODS). On the ‘Tools’ page in FluNexus, users can perform online analyses by uploading HA1 sequences and specifying the desired computational tool, virus subtype, and types of antigenic distance.

### 2.3 Characterization of benchmark datasets

The benchmark datasets are composed of paired HA1 sequences and HI titers. We focus on the HA1 region of the HA protein, as amino acid substitutions within this region—particularly those occurring at or near antigenic sites—play a pivotal role in driving immune escape and antigenic drift [7, 25, 26]. The HA1 datasets comprise 6,727 sequences of 326 amino acids for H1 (1976–2024), 6,353 sequences of 328 amino acids for H3 (1968–2024), and 77 sequences of 317 amino acids for H5 (1996–2020). Within these HA1 regions of each subtype, variable amino acid sites (Figure 2A), defined as those with the most frequent residue in less than 75% of sequences, often correspond to known antigenic epitopes. Substitutions at these critical sites, such as sites 156, 164, and 186 in H1 [27–29]; 156 and 193 in H3 [30, 31]; and 185 and 189 in H5 [32, 33], are recognized as primary drivers of antigenic drift.

**FIGURE 2.**
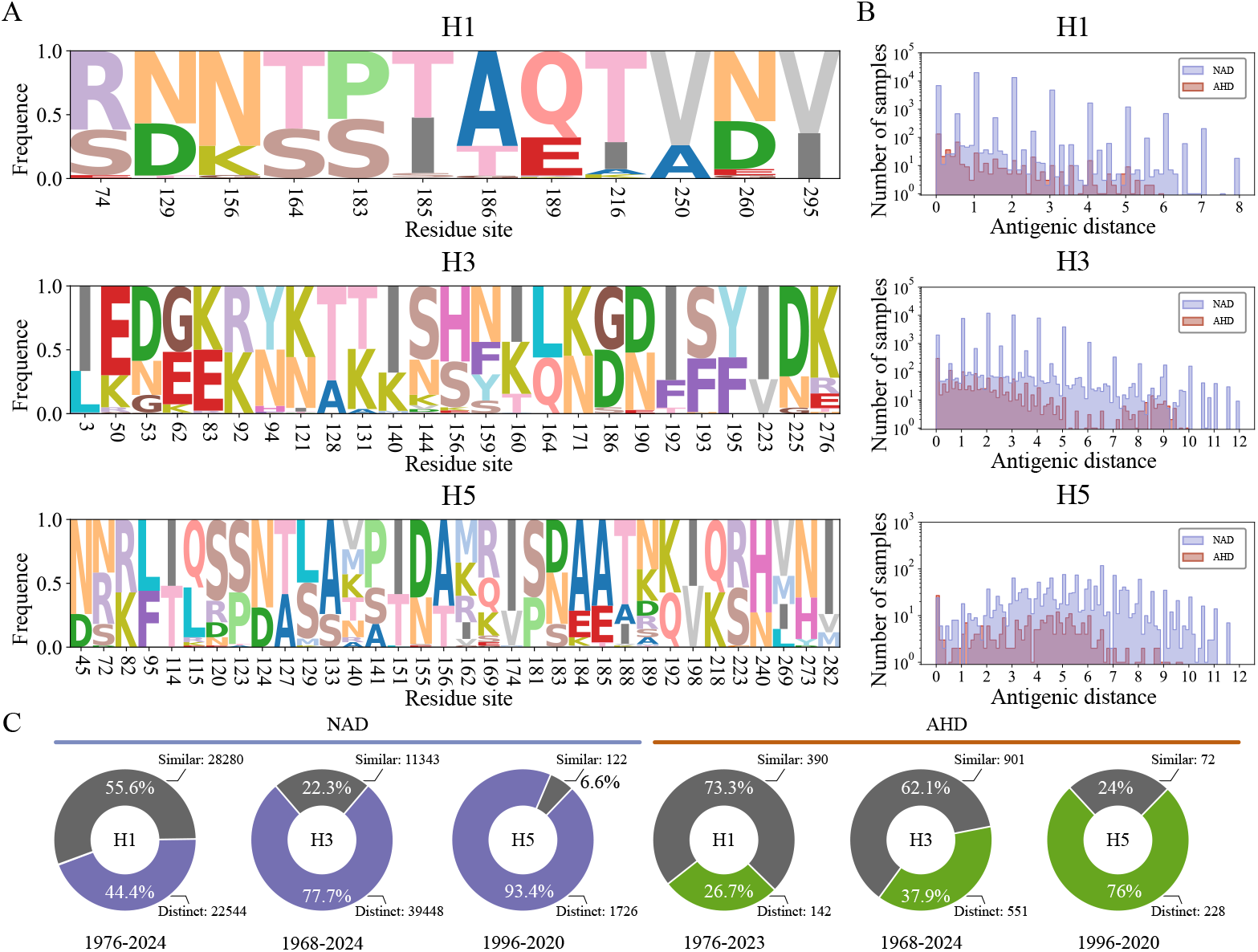
Overview of HI and HA1 datasets for H1, H3, and H5 influenza A sub-types. **(A)** Sequence logo plots of mutation hotspot regions within HA1 for each subtype. **(B)** Distribution of antigenic distances. **(C)** Distribution of the number of antigenically similar and distinct samples.

HI data are quantitative measurements derived from the HI assay, which provides the assessments of antigenicity among influenza virus isolates within the same subtype. The assay involves serial dilution of serum samples followed by incubation with influenza viruses and red blood cells to measure the ability of antibodies to inhibit HA-mediated agglutination [34]. In this study, we compile comprehensive HI benchmark datasets for H1, H3, and H5 influenza A subtypes. H1 includes 6,727 antigens and 79 antisera collected between 1976 and 2024, while H3 comprises 6,329 antigens and 243 antisera spanning from 1968 to 2024. For the H5 subtype, the collection contains 77 antigens and 24 antisera covering the period from 1996 to 2020. Regarding data sources, HI data for the H1 subtype are exclusively retrieved from the annual and interim reports of the Worldwide Influenza Centre at the Francis Crick Institute. For the H3 subtype, we integrate data sourced from the Worldwide Influenza Centre at the Francis Crick Institute with the dataset from Smith et al.[7], which comprises 79 vaccine strains and 253 reference viruses spanning the period from 1968 to 2003. Meanwhile, we generate a comprehensive H5 subtype–specific HI dataset in our BSL-3 laboratory. This dataset spans most globally reported H5 antigenic groups, thereby bridging critical gaps in existing data resources. Specifically, antisera are raised in three specific-pathogen-free (SPF) chickens immunized with 0.5 mL of an inactivated vaccine virus carrying the indicated HA and NA genes, with six remaining internal genes derived from A/Puerto Rico/8/1934 (PR8), as described previously [35]. Sera from the three chickens are pooled and stored at -80 °C for subsequent antigenic analyses. HI assays are performed according to the WHO-recommended protocol [36].

### 2.4 Benchmarking of antigenic prediction methods

We present a comprehensive benchmarking study to evaluate the performance of antigenic prediction methods, aiming to elucidate the strengths and limitations of distinct algorithms in practical scenarios and facilitate rational model selection tailored to specific subtypes and research inquiries. All ten methods are implemented and tested within a unified framework using identical datasets and evaluation metrics. The benchmarking framework is developed based on three key criteria: (i) the ability to discriminate antigenic variants from non-variants, measures by the Area Under the Receiver Operating Characteristic Curve (AUC) and Area Under the Precision-Recall Curve (AUPRC), (ii) the accuracy in inferring antigenic distances between virus pairs, measured by the Mean Absolute Error (MAE) and Mean Squared Error (MSE), and (iii) computational efficiency, quantified by the time for HA1 sequence encoding and model training. To ensure robust evaluation, we employ both crossvalidation and retrospective data partitioning strategies (see Supplementary S2 for details). Ultimately, we synthesize these results into a comprehensive performance summary (Figure 3D, Supplementary Figure S3).

**FIGURE 3.**
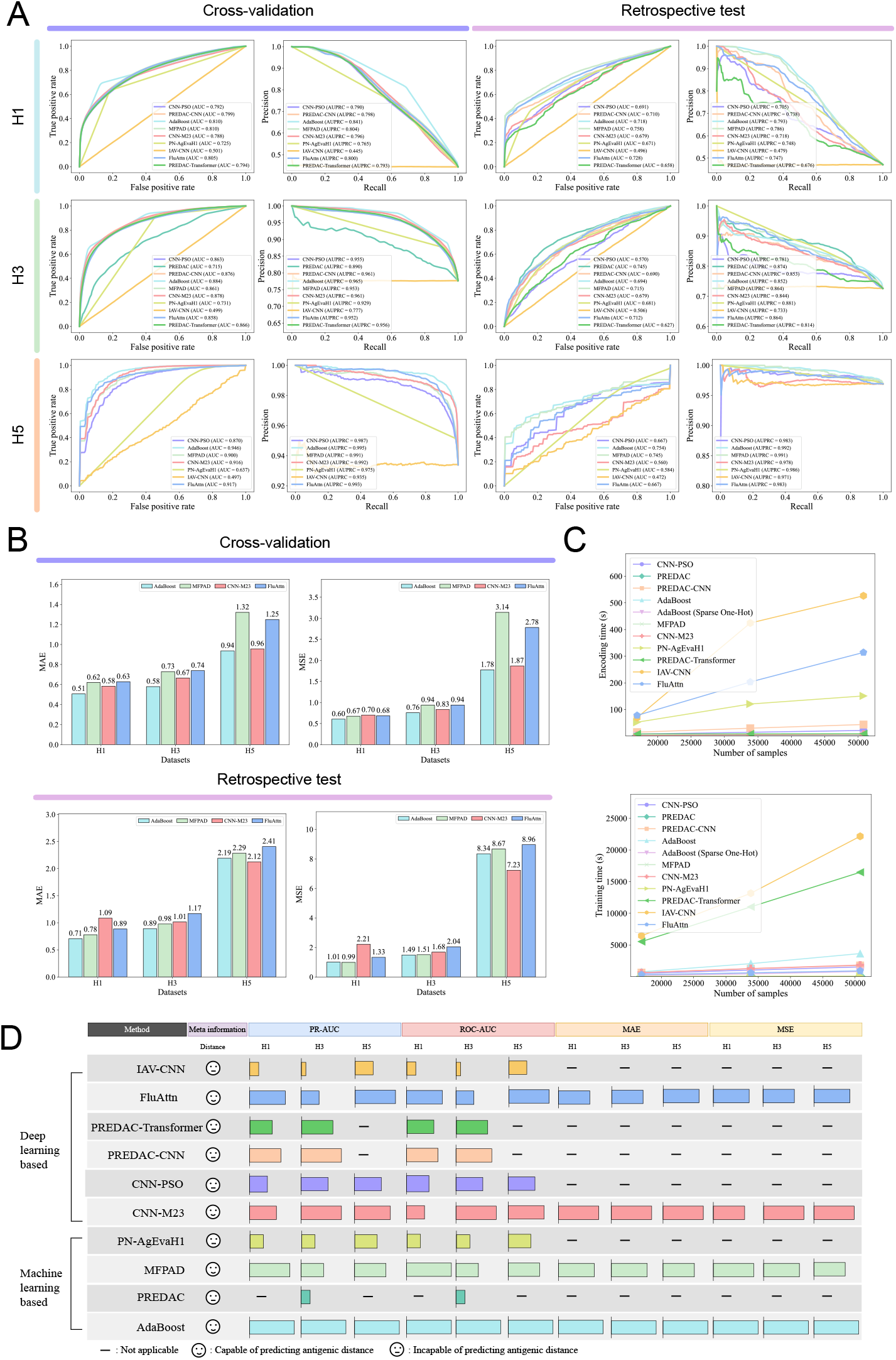
Performance evaluation of methods using the NAD antigenic distance.**(A)** Performance of discriminating antigenic variants from non-variants. **(B)** Performance of inferring antigenic distance. **(C)** Comparison of feature encoding and training time across methods for H3 subtype. **(D)** Summary of method performance. The bars represent the relative rankings of model performance, with longer bars indicating better performance.

In experiments of discriminating antigenic variants from non-variants using NAD antigenic distance (Figure 3A), AdaBoost exhibits superior performance across H1, H3, and H5 subtypes under the cross-validation, while FluAttn, MFPAD, and AdaBoost maintain high predictive accuracy in the retrospective testing scenario. For AHD-based classification, MFPAD, AdaBoost, and FluAttn prove superior under both the cross-validation and retrospective testing strategies (Supplementary Figure S1A). Notably, we observe a performance decline for nearly all models under retrospective testing, suggesting a constrained capacity to generalize to temporally distant or unseen viral strains.

Regarding antigenic distance inference between virus pairs based on NAD antigenic distance (Figure 3B), AdaBoost and CNN-M23 consistently yield optimal performance across H1, H3, and H5 subtypes under cross-validation. AdaBoost further exhibits exceptional efficacy in retrospective testing, especially for the H3 subtype. In terms of the AHD antigenic distance (Supplementary Figure S1B), MFPAD demonstrates robust predictive accuracy, specifically for the H1 and H5 subtypes. It is also evident that retrospective testing results in increased prediction errors relative to cross-validation across all subtypes, with the most significant increase occurring in H5 subtype.

Computational efficiency is assessed by measuring HA1 sequence encoding and model training time at incrementally increasing data scales. Observing that the original one-hot encoding in AdaBoost creates a dimensionality bottleneck, we implement a sparse vector representation, denoted AdaBoost (Sparse One-Hot), to mitigate this overhead. All benchmarking is conducted using 32-thread parallelization. As illustrated in Figure 3C and Supplementary Figure S2, MFPAD and PREDAC achieve the highest encoding efficiency, while MFPAD, PN-AgEvaH1, and PREDAC demonstrate superior efficiency in model training time. Conversely, IAV-CNN and FluAttn suffer from high encoding cost, attributed respectively to the computationally intensive trigram sliding-window process and the requirement for dynamic feature mining. IAV-CNN also presents the highest training computational cost owing to its dense matrix operations. Notably, for the H1 subtype (NAD distance), AdaBoost displays significantly elevated training times, highlighting its limited scalability when applied to datasets containing large numbers of unique antigens and antisera.

### 2.5 Improved mapping of antigenic evolution and visualization

HI data are typically generated by testing a limited number of antisera against a panel of viral isolates, resulting in numerous missing antigen-antiserum titers. This data sparsity poses challenges for the widely-used antigenic visualization tool Racmacs, hindering the accurate depiction of antigenic relationships among large sets of viruses (Figure 4). To improve the accuracy of the antigenic map, we propose a novel manifold-based algorithm (see METHODS) to more accurately position antigens and antisera, learning the underlying antigenic characterization in HI data. We employ the HI dataset to construct antigenic maps, and conduct a comparative benchmark against Racmacs (https://github.com/acorg/Racmacs) [7], Uniform Manifold Approximation and Projection (UMAP) [37], principal component analysis (PCA) [38], and t-distributed Stochastic Neighbor Embedding (t-SNE) [39] (see METHODS).

**FIGURE 4.**
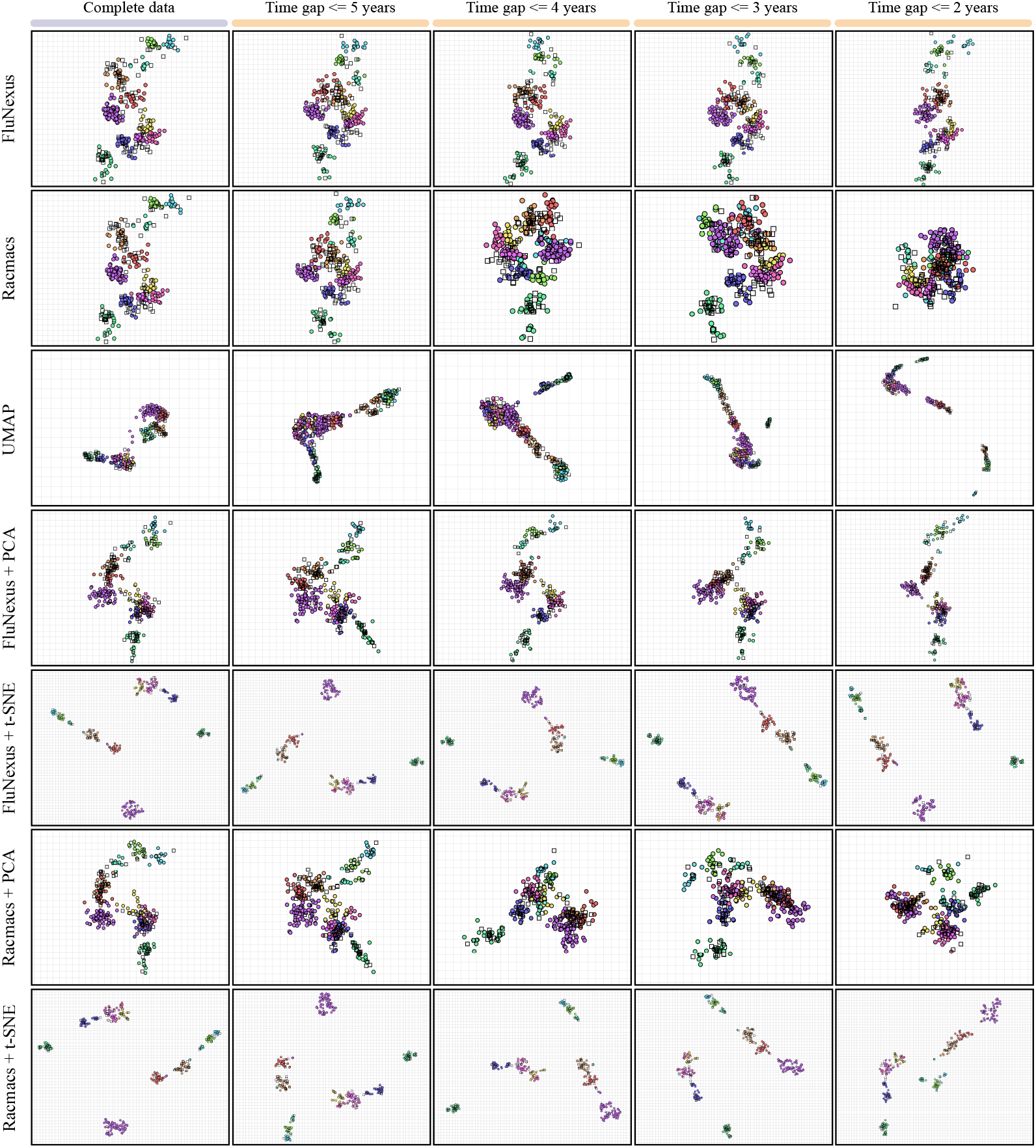
Antigenic maps generated by FluNexus, Racmacs, UMAP, PCA, and t-SNE using the HI data of H3 subtype sourced from Smith et al. under temporal sampling. Antigens in maps are colored consistently with the Smith et al. study.

To assess the capacity of FluNexus in preserving global evolutionary patterns under data-limited conditions, we simulate data scarcity by progressively excluding HI measurements where the antigen-serum temporal interval exceeds thresholds of 5, 4, 3, and 2 years. This simulation strategy aligns with real-world practices, where HI assays typically prioritize antigen-serum pairs isolated within close temporal proximity. We first benchmark the proposed method against established antigenic mapping tools using the HI data for H3 subtype from Smith et al. [7], a widely canonical reference for antigenic evolution. The results demonstrate that FluNexus outperforms other methods (Figure 4) by preserving coherent antigenic evolutionary trajectories and exhibiting robustness under both complete and temporally restricted data conditions. Furthermore, quantitative analysis confirms the superior stability of FluNexus, which achieves the highest concordance with true antigenic relationship from high-dimensional immunological data (MAE=0.64, MSE=0.84 for complete HI titers, Supplementary S3, Supplementary Tables S1 and S2). This suggests that our approach robustly captures the antigenic relationships from high-dimensional immunological data, effectively mitigating the noise associated with temporal sampling. Thus, the FluNexus-derived antigenic maps can be reliably applied to detect antigenic variants and prioritize vaccine strains in response to rapid viral evolution.

We subsequently evaluate FluNexus using independent HI datasets curated from the annual and interim reports of the Worldwide Influenza Centre at the Francis Crick Institute (2003–2024). In contrast to the Smith dataset, which possesses a canonical reference antigenic map, the post-2003 H3 evolutionary landscape lacks a unified, widely accepted expert-curated standard. To ensure robust map identifiability in this context, we restrict the analysis to viruses that present as both test antigens and reference antisera. The results show that FluNexus generates stable antigenic maps that preserve interpretable global evolutionary patterns and remains robust against data sparsity induced by temporal sampling (Supplementary Figure S4). Extended evaluation on H1 and H5 datasets also validates the subtype generalizability of FluNexus (Supplementary Figure S5 and S6).

The FluNexus web platform incorporates two interactive visualization modules: the antigenic map and the antigenic cluster. Users can construct antigenic maps using both the proposed manifold-based method and the established Racmacs method, with support for both HI data and NAD value inputs. An integrated control panel facilitates the refinement of the generated map, allowing for the adjustment of key parameters such as cartographic dimensionality (projecting antigens and antisera in high-dimensional immunological data to 2D or 3D cartography), the number of clusters, the random seed and the number of optimizers. Furthermore, point properties (e.g., size, shape, and color) remain dynamically adjustable post-rendering to enable customized visualization. The antigenic cluster module in FluNexus is implemented in accordance with PREDAC [10] (see METHODS for details). The initial positions of the points are determined using a built-in ‘spring’ layout (see Supplementary S4 for details). Similar to the antigenic map, users can adjust the size, shape, and color of points, and manually refine the layout via drag-and-drop.

## 3 CONCLUSION

In this study, we present FluNexus, the first-of-its-kind, one-stop-shop web platform that streamlines antigenic prediction and visualization for influenza A viruses. FluNexus is a versatile platform that integrates interactive modules for data preprocessing, online antigenic prediction with practical guidance for researchers, and visualization of influenza A virus antigenic evolution. Specifically, FluNexus proposes a novel manifold-based method for positioning antigens and antisera, ensuring the generation of more accurate antigenic cartographies, particularly when HI data is sparse.

FluNexus lowers the technical barrier, empowering researchers without specialized computational backgrounds to perform antigenic analysis of influenza A virus. Notably, beyond its application on influenza A virus, the core workflow and visualization framework of FluNexus can also be applied to other pathogens when assay-derived antigenic measurements and corresponding genomic sequences are available. Overall, FluNexus is poised to significantly aid virologists and public health officials in tracking antigenic evolution and improving vaccine strain selection.

## 4 METHODS

### 4.1 Data collection and preprocessing

We obtain all HA protein sequences of influenza A virus subtypes H1 and H3 from the GISAID database (https://gisaid.org/) [40], and additionally incorporate HA sequences of H3 subtype from Smith et al. [7]. For the H5 subtype, HA sequences are retrieved from GISAID and supplemented with additional sequences derived from virus isolates collected in China and processed in our BSL-3 laboratory. Sequences for each subtype are aligned separately using MAFFT, with A/California/04/2009 (H1N1) (isolate ID: EPI_ISL_376192), A/Aichi/2/1968 (H3N2) (isolate ID: EPI_ISL_123225) and A/Vietnam/1203/2004 (H5N1) (isolate ID: EPI_ISL_10656749), serving as reference strains for H1, H3 and H5, respectively. After alignment, only the HA1 subunit is retained, and the signal peptides (17, 16, and 16 amino acids for H1, H3, and H5 viruses, respectively) are removed. Sequences containing ambiguous amino acid codes (“B”, “Z”, “J”, or “X”) or with more than 10% gaps are excluded. Additionally, sequences with gaps at both ends are discarded, as these typically represent incomplete sequencing artifacts rather than true biological variations.

We collect HI titer data for H1 and H3 subtypes from 43 semiannual reports published by the Worldwide Influenza Centre at the Francis Crick Institute (https://www.crick.ac.uk/). Meanwhile, we further integrate HI measurements of H3 subtype for 79 vaccine strains and 253 reference viruses reported by Smith et al. [7] from 1968 to 2003. H5 subtype-specific HI titers are obtained from parallel HI assays performed in a BSL-3 laboratory, using representative Chinese H5 isolates and a serum panel of 27 samples covering 17 phylogenetic clades. The raw data are converted from a non-editable format into a standardized, unified data format to ensure compatibility with computational tools and analytical workflows. Meanwhile, the non-numeric titer values in the HI data are converted into numeric values following the procedure of Smith et al. [7].

The antigen-antiserum distances in the antigenic map corresponding to HI values are quantified using two measurements, NAD and AHD, introduced in previous studies [15, 24, 41]. Let *H*_*i,j*_ denote the HI titer measured against antigen *i* using antiserum from strain *j*. NAD can be defined as:

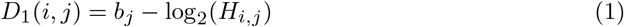

where *b*_*j*_ represents the log_2_ of the maximum HI titer observed across all antigens for antiserum *j*. A difference of two or more units in NAD between two viruses generally indicates a significant antigenic drift [7], reflecting antigenic changes sufficient to reduce protective immunity and prompting vaccine updates. AHD is defined as:

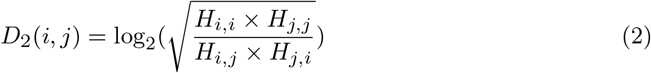

where *H*_*j,i*_ denotes the HI titer measured using antiserum derived from strain *i* against antigen *j, H*_*i,i*_ denotes the homologous titer measured using antiserum from strain *i* against its own antigen, and likewise for *H*_*j,j*_ . Log_2_ transformation can compress the value distribution, alleviate data sparsity, and improve computational stability [21]. When the AHD distance between two viruses reaches or exceeds 2, they are considered antigenically distinct [17, 21, 23, 24].

### 4.2 Settings of benchmarked methods

All machine learning methods are implemented following their studies. For deep learningbased methods, including CNN-PSO, CNN-M23, PREDAC-CNN, IAV-CNN, PREDAC-Transformer, and FluAttn, we evaluate the performance of models on the test set using the optimal checkpoint corresponding to the lowest loss on the validation set across 1,000 training epochs.

The basic principles of the ten different methods are elucidated as follows:

- **PREDAC**. PREDAC employs a Naive Bayes classifier to discriminate antigenic variants from non-variants by analyzing amino acid substitutions in HA1 sequences. It integrates diverse structural and physicochemical features, including mutations in epitopes, receptor-binding sites, and glycosylation substitutions, alongside shifts in physicochemical properties.
- **CNN-PSO**. CNN-PSO utilizes particle swarm optimization (PSO) to simultaneously select optimal physicochemical features and refine the architecture of a convolutional neural network (CNN), thereby achieving high-accuracy predictions of influenza virus antigenicity.
- **CNN-M23**. CNN-M23 encodes HA1 sequences by applying PCA to the full AAindex, reducing amino acid representations to 11-dimensional vectors. This method transforms sequences into structured numerical matrices and employs an AlexNet-inspired CNN to predict pairwise antigenic distances, effectively capturing the complex mutation patterns driving antigenic drift.
- **MFPAD**. MFPAD integrates multidimensional feature representations of HA1 sequences—comprising mutation counts, key antigenic sites, epitopes, glycosylation sites, and physicochemical properties—to train an XGBoost model for robust antigenicity prediction.
- **PREDAC-CNN**. REDAC-CNN constructs subtype-specific feature dictionaries for H1N1 and H3N2 derived from physicochemical properties in the AAindex. By encoding HA1 sequences with these dictionaries, it utilizes a CNN to capture local amino acid interactions, demonstrating superior performance in variant identification.
- **AdaBoost**. AdaBoost combines HA1 sequence data with associated metadata to predict antigenic distances. It leverages an adaptive boosting algorithm to learn a seasonally updated mapping from genetic variations to antigenic changes.
- **IAV-CNN**. IAV-CNN utilizes ProtVec to map amino acid 3-grams into 100-dimensional vectors, effectively capturing strain variability. It features a 2D CNN augmented with Squeeze-and-Excitation (SE) blocks to model channel interdependencies, thereby focusing the network’s attention on residues critical for antigenicity.
- **PN-AgEvaH1**. PN-AgEvaH1 employs a random forest framework to evaluate antigenicity by integrating physicochemical properties with structure-derived correlations, accurately mapping sequence variations to antigenic characteristics.
- **PREDAC-Transformer**. PREDAC-Transformer leverages a transformer encoder with self-attention mechanisms to capture long-range sequence dependencies. It synergizes physicochemical properties with ESM-2 protein language model embeddings to enhance antigenic prediction capabilities.
- **FluAttn**. FluAttn introduces an attention-based feature mining framework to automatically identify and weight antigenicity-relevant properties from the AAindex. These adaptive features are combined with sequence differences to train a multilayer perceptron for precise antigenic prediction.

### 4.3 Improved antigenic map

The shape space theory [42] shows that antigen-antiserum distance metrics can be transformed into meaningful spatial representations, allowing antigenic relationships to be visualized in the form of antigenic maps. Smith et al. [7] have employed MDS, based on the NAD values, to embed both antigens and antisera into a low-dimensional Euclidean space, thereby enabling the construction of an antigenic map. However, the NAD matrix is characteristically sparse. This sparsity is inherent to the temporal bias [8] and can introduce the unrealistic positioning of antigenically distant strains in close proximity on the antigenic map.

Manifold-based methods offer an effective way to capture the intrinsic geometry of complicated data, addressing limitations of linear methods such as MDS when Euclidean distances fail to reflect true relationships. By incorporating geodesic information that reflects the underlying manifold structure, Isomap [43] successfully uses locally reliable measurements to recover the global structure of the underlying data space. Consequently, inspired by Isomap, we introduce geodesic distance to estimate the distance for antigen–antigen, antiserum–antiserum, and antigen–antiserum pairs that lack HI measurements. We first construct a weighted antigen–antiserum graph where the nodes represent the antigens or antisera and the edges are weighted by the NAD distance. This initial graph is then refined into a neighborhood graph by connecting each antigen or antiserum to its *k* nearest neighbors. Geodesic distances on this refined graph are then calculated using Dijkstra’s algorithm. Then, low-dimensional embedding will be constructed based on the NAD and geodesic distances and the Limited-memory Broyden–Fletcher–Goldfarb–Shanno algorithm[44] is adopted to optimize the low-dimensional coordinates of antigens and antisera. Therefore, FluNexus can best preserve the mainfold’s estimated intrinsic geometry and ensure the generation of more accurate antigenic cartographies. Details can be found in Supplementary S5.

### 4.4 Settings of antigenic visulization

With the exception of PCA and t-SNE, all employed methods map antigens and antisera directly into a two-dimensional space. However, standard PCA and t-SNE implementations are sensitive to missing entries inherent in HI data. To address this limitation, we adopt a two-stage dimensionality reduction strategy for these two methods. First, we embed antigens and antisera into a 5-dimensional space using either FluNexus or Racmacs [7] to impute missing information. Subsequently, PCA or t-SNE is applied to map these high-dimensional embeddings into a 2-dimensional plane.

Following dimensionality reduction, k-means clustering is performed on the antigens. The number of clusters (*k*) is determined specifically for each dataset: H1 (*k* = 4), H3 derived from Smith et al.[7] (*k* = 11), H3 collected from WIC (*k* = 13), and H5 (*k* = 18). Both FluNexus and Racmacs are configured with 20,000 optimization runs and a fixed random seed of 0 to ensure reproducibility.

The fundamental principles and parameter configurations of the comparative antigenic mapping methods are detailed as follows:

- **Racmacs**. Racmacs employs a modified MDS algorithm to position antigens and antisera within a low-dimensional Euclidean space. The algorithm optimizes coordinates to minimize the error between map-derived distances and experimental distances. In this study, Racmacs is implemented following the protocols of Smith et al.[7] using default settings.
- **UMAP**. UMAP constructs a fuzzy topological representation of the dataset by approximating a manifold, assuming the data is uniformly distributed on a locally connected Riemannian manifold. The algorithm optimizes the low-dimensional embedding to minimize the fuzzy set cross-entropy between the high-dimensional and low-dimensional topological structures, thereby preserving both local and global structural features. The hyperparameters for UMAP are set as follows: `n_neighbors` = 15, `metric` = ‘euclidean’, `init` = ‘spectral’, and `min_dist` = 0.1.
- **PCA**. PCA reduces dimensionality by linearly mapping high-dimensional data into a lower-dimensional subspace spanned by orthogonal principal components that maximize the retained variance. In this study, PCA is implemented using *scikit-learn* with default settings, projecting data onto the first two principal components.
- **t-SNE**. t-SNE quantifies high-dimensional similarities as conditional probabilities and models low-dimensional embeddings using a heavy-tailed Student’s t-distribution to mitigate the crowding problem. The algorithm minimizes the Kullback-Leibler divergence between these distributions to effectively preserve local manifold structures. Specifically, we employ the *scikit-learn* implementation with the following hyperparameters: `perplexity` = 30, learning rate = ‘auto’, `metric` = ‘euclidean’, and `max_iter` = 1,000.

In the interactive visualization, antigenic clustering follows the PREDAC methodology using the Markov-clustering Python package with default parameters.

## 5 AUTHOR CONTRIBUTIONS

**Xingyi Li**: Conceptualization; funding acquisition; investigation; project administration; supervision; visualization; writing—review and editing. **Chunyan Zhou**: Data curation;methodology; software; writing—original draft. **Han Wu**: Data curation; methodology; software; writing—original draft. **Kexin Xiao**: Formal analysis; investigation; validation. **Jun Hao**: Formal analysis; investigation; validation. **Dongmin Zhao**: Formal analysis; investigation; validation. **Junnan Zhu**: Formal analysis; investigation; validation. **Yue Li**: Software; validation. **Jiajie Peng**: Software; validation. **Jia Gu**: Software; validation. **Guohua Deng**: Data curation; formal analysis. **Weigang Cai**: Data curation; formal analysis. **Min Li**: Writing—review and editing; validation; supervision; project administration. **Yan Liu**: Data curation; formal analysis; supervision; project administration. **Xuequn Shang**: Resources; methodology; funding acquisition; project administration. **Hualan Chen**: Data curation; resources; methodology; supervision; project administration. **Huihui Kong**: Data curation; formal analysis; methodology; supervision; funding acquisition; project administration; writing—review and editing.

## 6 ACKNOWLEDGMENTS

This work is supported in part by the National Key Research and Development Program of China [2022YFD1801200], the Agricultural Science and Technology Innovation Program [CAAS-CSLPDCP-202301], the State Key Laboratory for Animal Disease Control and Prevention Foundation [SKLADCPKFKT202407], the Macau Young Scholars Program [AM2024027], the Young Talent Fund of Xi’an Association for Science and Technology [0959202513204x]. We acknowledge the use of HI data in the annual and interim reports generated by the Worldwide Influenza Centre at the Francis Crick Institute. We also acknowledge the use of HA sequence data obtained from the GISAID database.

## 8 DATA AVAILABILITY

HA1 sequences for H1 and H3 influenza viruses can be downloaded from GISAID[40] at https://gisaid.org/. Corresponding HI data for H1 and H3 subtypes can be downloaded at https://www.crick.ac.uk/research/platforms-and-facilities/worldwide-influenza-centre/annual-and-interim-reports. Additional HA1 sequences and HI data of H3 subtype can be downloaded from the study by Smith et al[7]. HA1 sequences and H5-specific HI data can be downloaded from GISAID and GitHub https://github.com/xingyili/FluNexus-methodbox/tree/main/H5 data, respectively. The source code implementing the ten antigenic prediction methods is available at https://github.com/xingyili/FluNexus-methodbox.

