## Supplementary for "FluNexus: a versatile web platform for antigenic prediction and visualization of influenza A viruses"

#### Supplementary Section

#### S1 Architecture and implementation of FluNexus

FluNexus adopts a decoupled architecture that separates the front-end and back-end components. The front end is responsible for rendering interactive interfaces and transmitting user requests to the back end, primarily implemented using HTML, CSS, and JavaScript. Antigenic map and antigenic cluster are visualized using Three.js (`https://threejs.org/`). Meanwhile, the back end orchestrates various computational tasks in response to user requests received from the front end and returns processed results to the client. It employs the Axum web framework (`https://github.com/tokio-rs/axum`) to handle HTTP request routing, processing, and manage WebSocket connections. In addition, FluNexus employs several strategies to safeguard user privacy. All data transmissions are secured via SSL certificates to ensure encrypted communication. No cookies are used for tracking or session management. Data submitted by the user resides only in memory for short-term processing and is never persisted on the server.

#### S2 Benchmark metrics, and partitioning strategies for benchmarking analysis

In this study, we evaluate the performance of each method using the following metrics: MAE and MSE for quantifying antigenic distance between virus pairs, AUC and AUPRC for evaluating the ability to distinguish antigenic variants from non-variants. All metrics are first computed separately for each individual fold or test year, and the overall performance is calculated as the average of the per-fold or per-year results, weighted according to the number of test samples in each fold or year.

For the experiment of distinguishing antigenic variants from non-variants, labels are assigned to virus pairs based on their antigenic relationship: pairs exhibiting antigenic variation are designated as positive samples, whereas pairs that are antigenically similar are designated as negative samples. Subsequently, AUC is computed as the area under the Receiver Operating Characteristic curve, which plots the True Positive Rate (TPR) against the False Positive Rate (FPR) across all decision thresholds, and quantifies the model's ability to discriminate between positive and negative virus pairs. Here,

$$\mathrm{TPR} = \frac{\mathrm{TP}}{\mathrm{TP} + \mathrm{FN}} \tag{1}$$

$$\mathrm{FPR} = \frac{\mathrm{FP}}{\mathrm{FP} + \mathrm{TN}} \tag{2}$$

where TP is the number of antigenically variant pairs correctly classified as variant, FP is the number of antigenically similar pairs incorrectly classified as variant, TN is the number of antigenically similar pairs correctly classified as similar, and FN is the number of antigenically variant pairs incorrectly classified as similar. AUPRC is computed as the area under the Precision-Recall curve, which plots Precision against Recall across different decision thresholds, and emphasizes performance on the positive class. Here,

$$\mathrm{Precision} = \frac{\mathrm{TP}}{\mathrm{TP} + \mathrm{FP}} \tag{3}$$

$$\mathrm{Recall} = \frac{\mathrm{TP}}{\mathrm{TP} + \mathrm{FN}} \tag{4}$$

For the antigenic distance prediction experiment, MAE and MSE can be calculated as follows:

$$\mathrm{MAE} = \frac{1}{n}\sum_{i=1}^{n} |y_i - \hat{y}_i| \tag{5}$$

$$\mathrm{MSE} = \frac{1}{n}\sum_{i=1}^{n} (y_i - \hat{y}_i)^2 \tag{6}$$

where $y_i$ denotes the true antigenic distance between the $i$-th pair of virus strains, $\hat{y}_i$ is the corresponding value predicted by the model, and $n$ is the total number of virus pairs (i.e., the total number of samples).

Finally, the overall performance across all folds or test years is then obtained by combining the per-fold or per-year results according to the number of test samples in each fold or year. Specifically, if $M_y$ denotes the metric for fold $y$ or year $y$, the overall metric is given by:

$$\mathrm{M}_{\text{average}} = \frac{\sum_{y=1}^{Y} N_y \, \mathrm{M}_y}{\sum_{y=1}^{Y} N_y} \tag{7}$$

where Y is the number of folds or test years and $N_y$ denotes the number of virus pairs in the test set for fold $y$ or year $y$.

During model evaluation, two data partitioning strategies are employed: cross-validation and retrospective testing. Cross-validation is conducted at the titer level using five-fold partitioning repeated ten times. For the retrospective testing, strains are partitioned according to their isolation year. Data collected prior to the target year are used for training and validation, while strains from the target year and the following year are reserved for testing. When a separate validation set is used, samples from the two years preceding the target year is designated for validation, and these validation samples are excluded from the training set. The target years for each dataset are listed below:

- NAD (H1 subtype): 2019, 2020, 2021, 2022, 2023
- NAD (H3 subtype): 2019, 2020, 2021, 2022, 2023
- NAD (H5 subtype): 2015, 2016, 2017, 2018, 2019
- AHD (H1 subtype): 2018, 2019, 2020, 2021, 2022
- AHD (H3 subtype): 2019, 2020, 2021, 2022, 2023
- AHD (H5 subtype): 2015, 2016, 2017, 2018, 2019

### S3 Quantitative metrics of antigenic map

To evaluate the accuracy of different antigenic mapping methods, we utilize MAE and MSE as performance metrics to quantify the deviation of the generated antigenic map from the original HI titer data. Following the study of Smith et al.[1], the MAE and MSE are defined as follows:

$$MAE = \frac{1}{|\mathcal{P}_{obs}|} \sum_{(i,j)\in\mathcal{P}_{obs}} \begin{cases} \sigma(D_{ij} - d_{ij}) \cdot |D_{ij} - d_{ij}|, & \text{if } (i,j) \in \mathcal{P}_{\text{obs}}^{\text{thr}} \\ |D_{ij} - d_{ij}|\,, & \text{otherwise} \end{cases} \tag{8}$$

$$MSE = \frac{1}{|\mathcal{P}_{obs}|} \sum_{(i,j)\in\mathcal{P}_{obs}} \begin{cases} \sigma(D_{ij} - d_{ij}) \cdot (D_{ij} - d_{ij})^2, & \text{if } (i,j) \in \mathcal{P}_{\text{obs}}^{\text{thr}} \\ (D_{ij} - d_{ij})^2\,, & \text{otherwise} \end{cases} \tag{9}$$

where $\mathcal{P}_{\text{obs}}$ denote the set of antigen-antiserum pairs. $\mathcal{P}_{\text{obs}}^{\text{thr}} \subset \mathcal{P}_{\text{obs}}$ contains antigen–antiserum pairs whose HI measurements are thresholded values, typically reported as being below a detection limit (e.g., $< 10$). $D_{ij}$ denotes the antigenic distance between antigen $i$ and antiserum $j$ derived from experimental HI titers, while $d_{ij}$ represents the Euclidean distance between the antigen $v_i$ and antiserum $v_j$ in the constructed antigenic map. The term $\sigma(\cdot)$ denotes a sigmoid function, introduced to modulate the penalty for thresholded values.

### S4 Layout algorithm for antigenic cluster visualization

To visualize the antigenic clusters, we employ Fruchterman-Reingold force-directed algorithm, which generates a force-directed layout by modeling edges as springs that draw connected nodes together, while treating all nodes as mutually repulsive entities, analogous to anti-gravity forces. The simulation iteratively updates node positions until a near-equilibrium state is reached. This layout provides a clear representation of the network structure, facilitating the visualization of both the global architecture and local organization of the antigenic clusters.

### S5 Improved antigenic map

Firstly, we represent each antigen and antiserum by a coordinate vector in a low-dimensional embedding space (e.g., 2D or 3D). Subsequently, let $D \in \mathbb{R}^{M\times N}$ denote the NAD matrix, where $M$ and $N$ represent the number of antigens and antisera, respectively. We then construct an undirected weighted graph $\mathcal{G} = (\mathcal{V}, \mathcal{E}, w)$, where the node set $\mathcal{V}$ contains all antigens and antisera, i.e., $\mathcal{V} = \{v_1, \ldots, v_{M+N}\}$. The first $M$ nodes correspond to antigens, and the remaining $N$ nodes correspond to antisera. An edge $(v_i, v_{M+j}) \in \mathcal{E}$ exists only if the NAD $D_{ij}$ is available, and the associated edge weight is assigned as $w(v_i, v_{M+j}) = D_{ij}$. To better preserve the local manifold structure, a K-Nearest Neighbors (KNN) graph is constructed by retaining undirected edges only between each node and its $k$ nearest neighbors based on the edge weights, then the geodesic distances between all pairs of nodes are calculated using Dijkstra's algorithm. This results in a symmetric geodesic distance matrix $\Delta \in \mathbb{R}^{(M+N)\times(M+N)}$, where each entry $\Delta_{ij}$ denotes the length of the shortest path between node $v_i$ and node $v_j$ in the KNN graph. Subsequently, in order to flexibly modulate the influence of geodesic distances in the embedding process, we apply an affine transformation to the original geodesic distances $\Delta_{ij}$:

$$\hat{\Delta}_{ij} = \alpha \cdot \Delta_{ij} + \beta \tag{10}$$

where $\alpha$ and $\beta$ are hyperparameters that control the scaling and offset of the geodesic distances. Based on the KNN graph and transformed geodesic distances, we define four disjoint sets of node pairs:

- $\mathcal{P}_{\text{obs}}$: the set of antigen–antiserum pairs with NADs measurements.
- $\mathcal{P}_{\text{ag-sr}}$: the set of antigen–antiserum pairs without NAD measurements. Pairs with geodesic distance $\hat{\Delta}_{ij} < r$ are excluded.
- $\mathcal{P}_{\text{ag-ag}}$: the set of antigen–antigen pairs. Pairs with geodesic distance $\hat{\Delta}_{ij} < r$ are excluded.
- $\mathcal{P}_{\text{sr-sr}}$: the set of antiserum–antiserum pairs. Pairs with geodesic distance $\hat{\Delta}_{ij} < r$ are excluded.

Here, $r > 0$ is a hyperparameter representing the minimum geodesic radius that excludes pairs with small geodesic distance in the high-dimensional graph. Subsequently, before defining the loss terms, we introduce a gating mechanism to softly activate penalties only when certain distance constraints are violated. Here,

$$\sigma(x) = \frac{1}{1 + \exp^{(-\eta x)}} \tag{11}$$

is a sigmoid function that smoothly activates the error only when $x > 0$, with $\eta > 0$ controlling the sharpness of the transition. With the above definitions, the optimization objective comprises the following four terms, designed to learn low-dimensional representations for all antigens and antisera such that the distances in the embedded space reflect the original NADs where available, and approximate the underlying manifold structure and the geodesic structure where direct measurements are missing.

(1) For pairs $(i, j) \in \mathcal{P}_{obs}$, we define the residual $\xi_{ij} = D_{ij} - d_{ij}$, where $d_{ij}$ denotes the Euclidean distance between nodes $v_i$ and $v_j$ in the low-dimensional embedding space, and minimize the squared difference between embedded and observed distances:

$$e_{obs} = \sum_{(i,j)\in\mathcal{P}_{obs}} \begin{cases} \sigma(\xi_{ij}) \cdot (\xi_{ij})^2, & \text{if } (i,j) \in \mathcal{P}^{\text{thr}}_{\text{obs}} \\ (\xi_{ij})^2, & \text{otherwise} \end{cases} \tag{12}$$

The subset $\mathcal{P}\text{obs}^{\text{thr}} \subset \mathcal{P}\text{obs}$ contains antigen–antiserum pairs whose HI measurements are thresholded values, typically reported as being below a detection limit (e.g., ¡10).

(2) For pairs $(i, j) \in \{\mathcal{P}_{ag-sr}, \mathcal{P}_{ag-ag}, \mathcal{P}_{sr-sr}\}$, we introduce a soft constraint that adds a penalty to the objective function only when $d_{ij} < \tilde{\Delta}_{ij}$, i.e., when the distance between the node pair is smaller than their corresponding geodesic distance:

$$e_k = \sum_{(i,j)\in\mathcal{P}_k} \sigma(\hat{\Delta}_{i,j} - d_{ij}) \cdot (\hat{\Delta}_{i,j} - d_{ij})^2 \quad k \in \{ag\text{-}sr,\ ag\text{-}ag,\ sr\text{-}sr\} \tag{13}$$

Finally, the total objective function is a weighted sum:

$$e = e_{obs} + \lambda_1 e_{ag-sr} + \lambda_2 e_{ag-ag} + \lambda_3 e_{sr-sr} \tag{14}$$

where $\lambda_1, \lambda_2, \lambda_3 \geq 0$ are hyperparameters that control the relative contribution of each term. We adopt the Limited-memory Broyden–Fletcher–Goldfarb–Shanno algorithm[2] to optimize the low-dimensional coordinates of antigens and antisera by minimizing the objective $e$.

Furthermore, based on the spatial distribution of antigens and antisera in the antigenic map, FluNexus provides a K-means clustering [3, 4] functionality to facilitate the identification of meaningful clusters. Subsequently, to determine a suitable number of clusters $k$, we use the average silhouette coefficient [5] across all samples to evaluate clustering performance for different values of $k$ and choose the value that maximizes the silhouette coefficient.

**Supplementary Table**

**Table S1.** Comparison of MAE for FluNexus and comparative methods on the HI data of H3 subtype derived from Smith et al. under temporal sampling.

| **Method** | **Complete data** | **Time gap ≤ 5** | **Time gap ≤ 4** | **Time gap ≤ 3** | **Time gap ≤ 2** |
|---|---|---|---|---|---|
| FluNexus | 0.64 | 0.82 | 0.78 | 0.88 | 0.99 |
| Racmacs | 0.65 | 0.85 | 1.15 | 1.21 | 1.62 |
| UMAP | 3.21 | 3.54 | 3.46 | 3.24 | 2.61 |
| FluNexus + PCA | 1.22 | 1.53 | 1.26 | 1.51 | 1.33 |
| FluNexus + t-SNE | 7.21 | 6.53 | 5.91 | 5.39 | 6.40 |
| Racmacs + PCA | 1.23 | 1.57 | 1.76 | 1.61 | 1.89 |
| Racmacs + t-SNE | 6.41 | 6.37 | 5.70 | 5.66 | 6.85 |

**Table S2.** Comparison of MSE for FluNexus and comparative methods on the HI data of H3 subtype derived from Smith et al. under temporal sampling.

| **Method** | **Complete data** | **Time gap ≤ 5** | **Time gap ≤ 4** | **Time gap ≤ 3** | **Time gap ≤ 2** |
|---|---|---|---|---|---|
| FluNexus | 0.84 | 1.63 | 1.50 | 1.86 | 2.37 |
| Racmacs | 0.84 | 1.78 | 3.39 | 3.77 | 6.19 |
| UMAP | 15.39 | 18.56 | 17.90 | 16.05 | 11.99 |
| FluNexus + PCA | 2.93 | 4.32 | 3.22 | 4.41 | 3.49 |
| FluNexus + t-SNE | 181.36 | 130.39 | 110.66 | 99.28 | 133.23 |
| Racmacs + PCA | 3.00 | 4.48 | 5.53 | 5.13 | 6.44 |
| Racmacs + t-SNE | 135.91 | 121.63 | 106.21 | 108.83 | 169.41 |

### Supplementary Figure

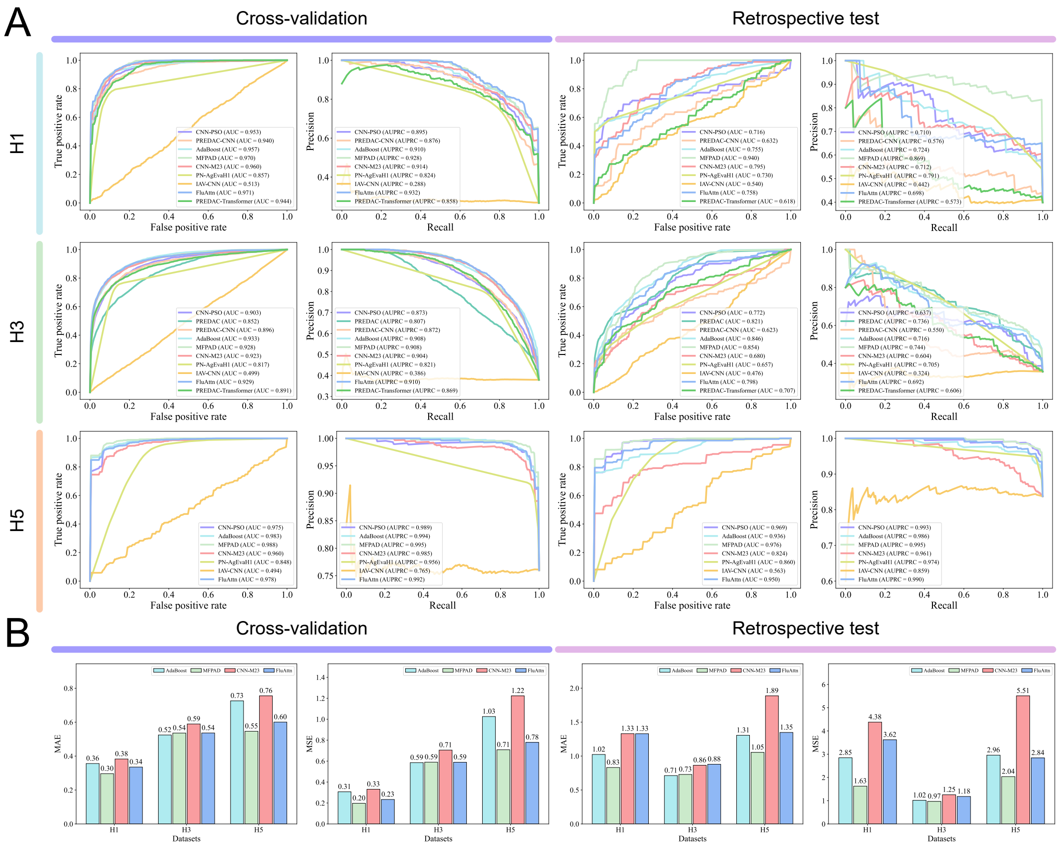


**Figure S1.** Performance evaluation of methods using the AHD antigenic distance. (A) Performance of different methods in distinguishing antigenic variants from non-variants. (B) Performance of different methods in inferring antigenic distance.

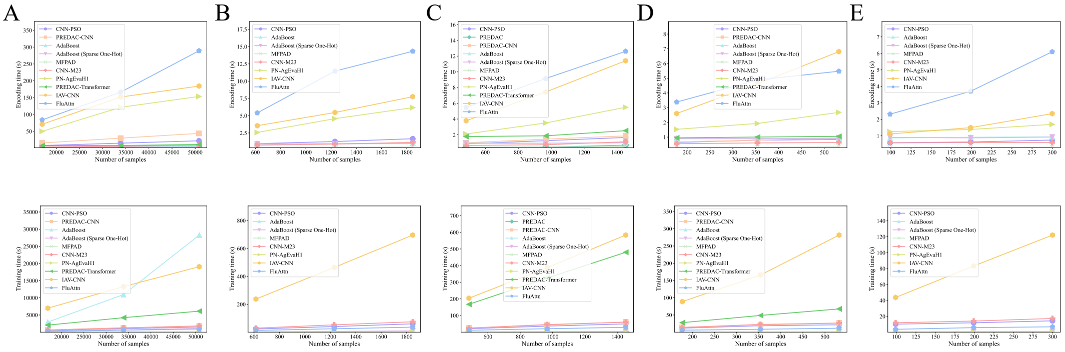


**Figure S2.** Comparison of feature encoding and training time across methods. **(A)** Feature encoding and training time across methods for H1 subtype using the NAD antigenic distance. **(B)** Feature encoding and training time across methods for H5 subtype using the NAD antigenic distance. **(C)** Feature encoding and training time across methods for H3 subtype using the AHD antigenic distance. **(D)** Feature encoding and training time across methods for H1 subtype using the AHD antigenic distance. **(E)** Feature encoding and training time across methods for H5 subtype using the AHD antigenic distance.

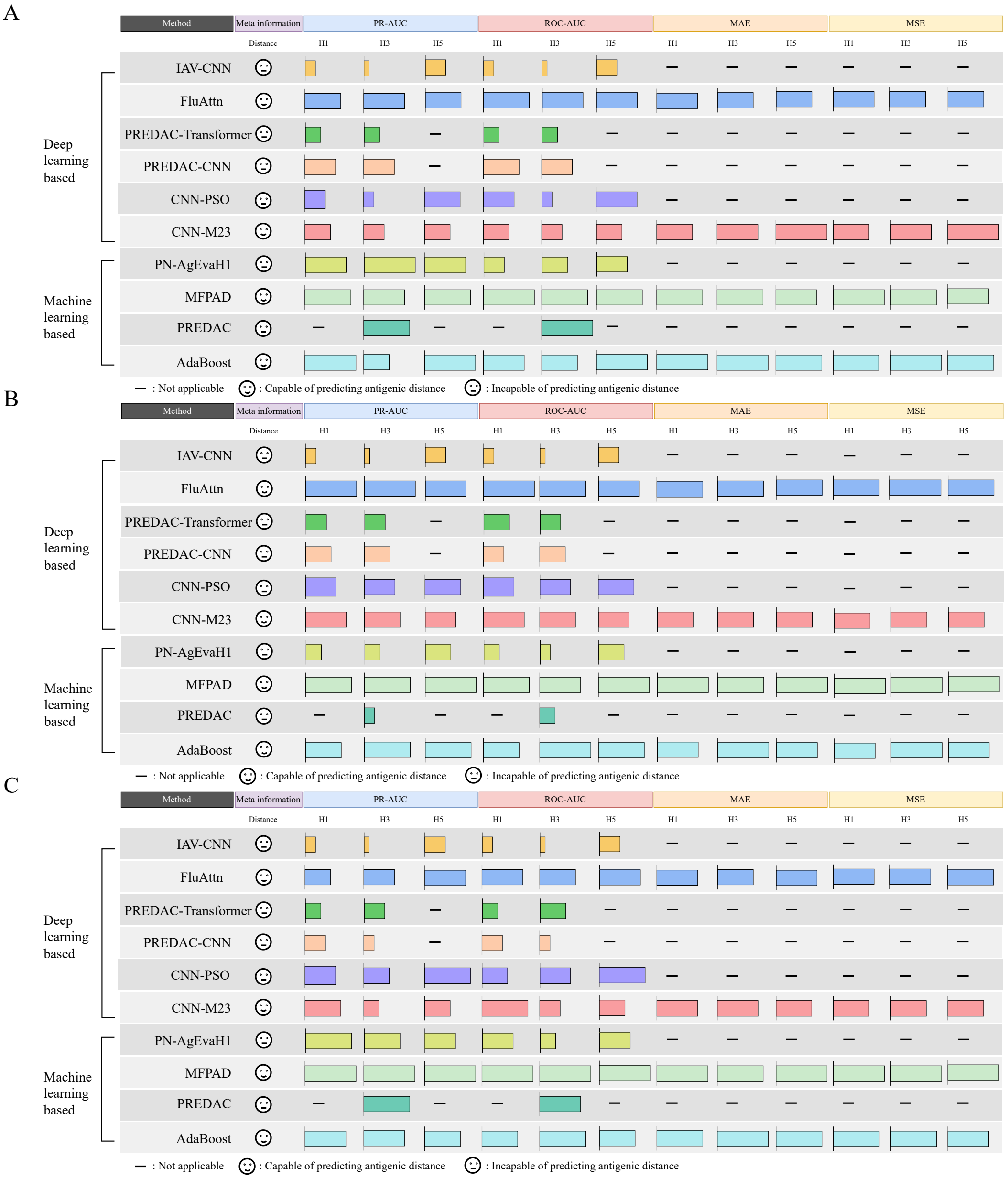


**Figure S3.** Overview of method performance. (A) Summary of method performance for the NAD distance metric under retrospective testing. (B) Summary of method performance for the AHD distance metric under cross-validation. (B) Summary of method performance for the AHD distance metric under retrospective testing.

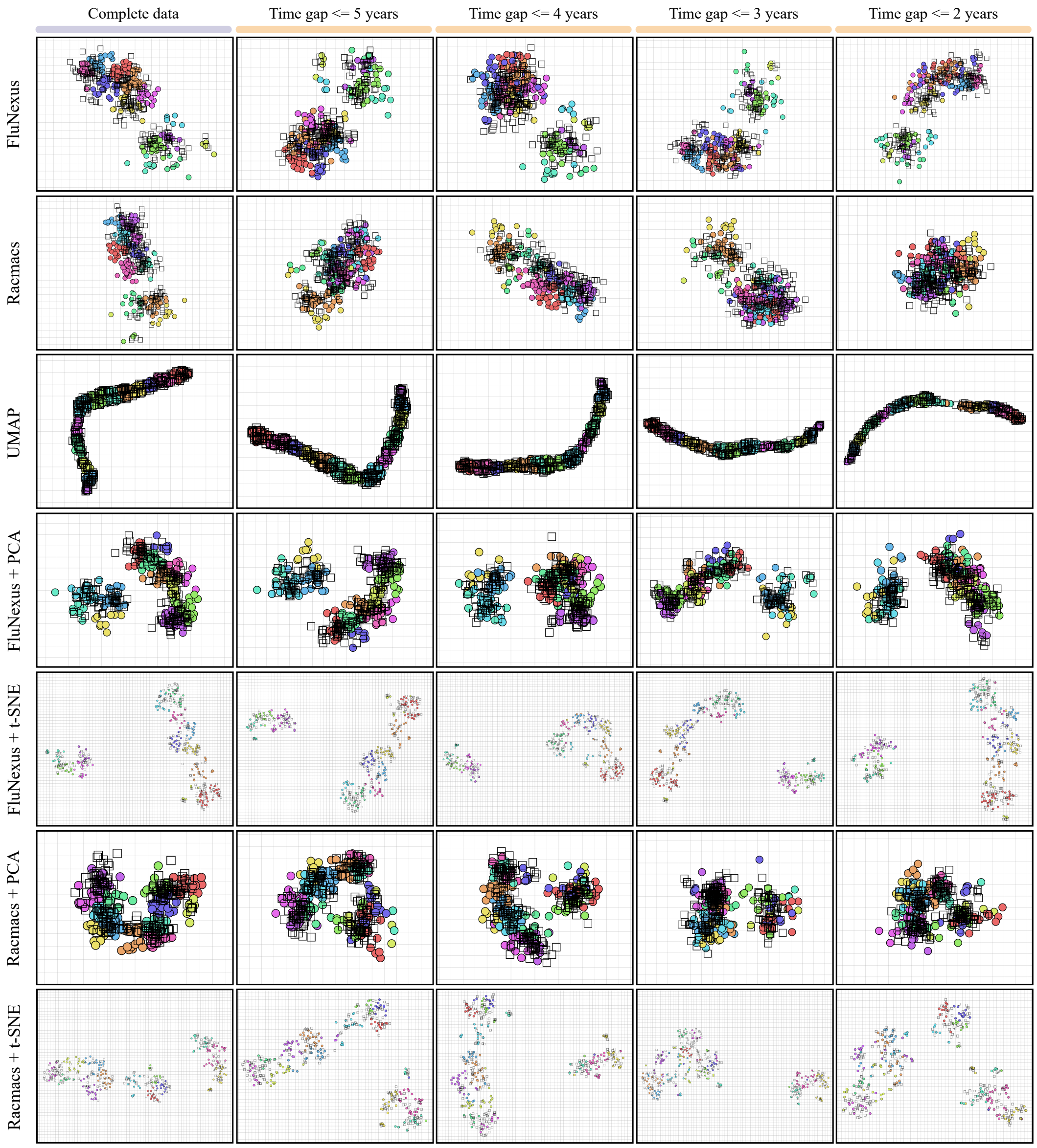


**Figure S4.** Antigenic maps generated by FluNexus, Racmacs, UMAP, PCA, and t-SNE using the HI data of H3 subtype derived from the annual and interim reports of the Worldwide Influenza Centre at the Francis Crick Institute under temporal sampling. For each method, antigens are colored consistent with antigenic map generated from their complete HI data.

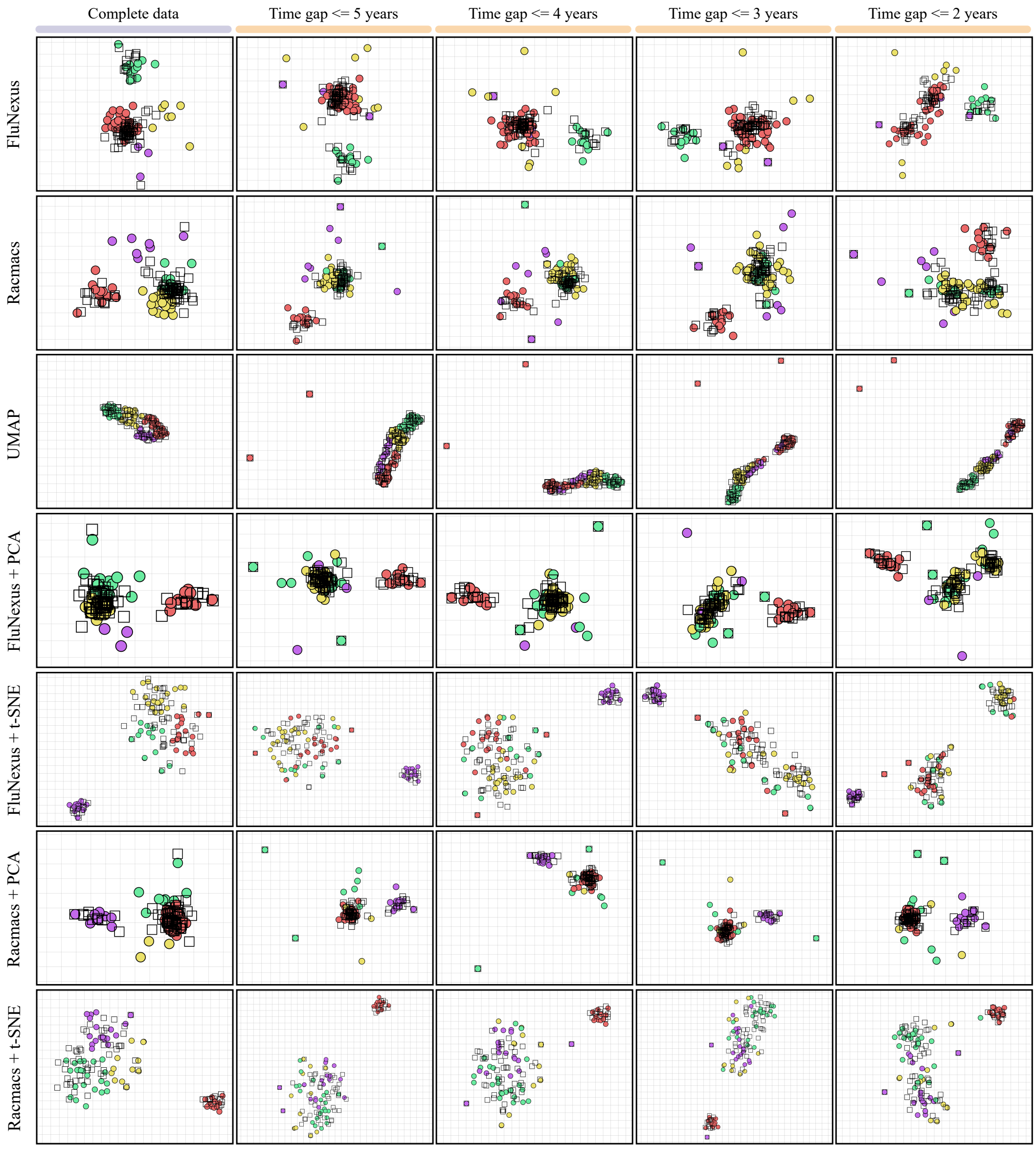


**Figure S5.** Antigenic maps generated by FluNexus, Racmacs, UMAP, PCA, and t-SNE using the HI data of H1 subtype under temporal sampling. For each method, antigens are colored consistent with antigenic map generated from their complete HI data.

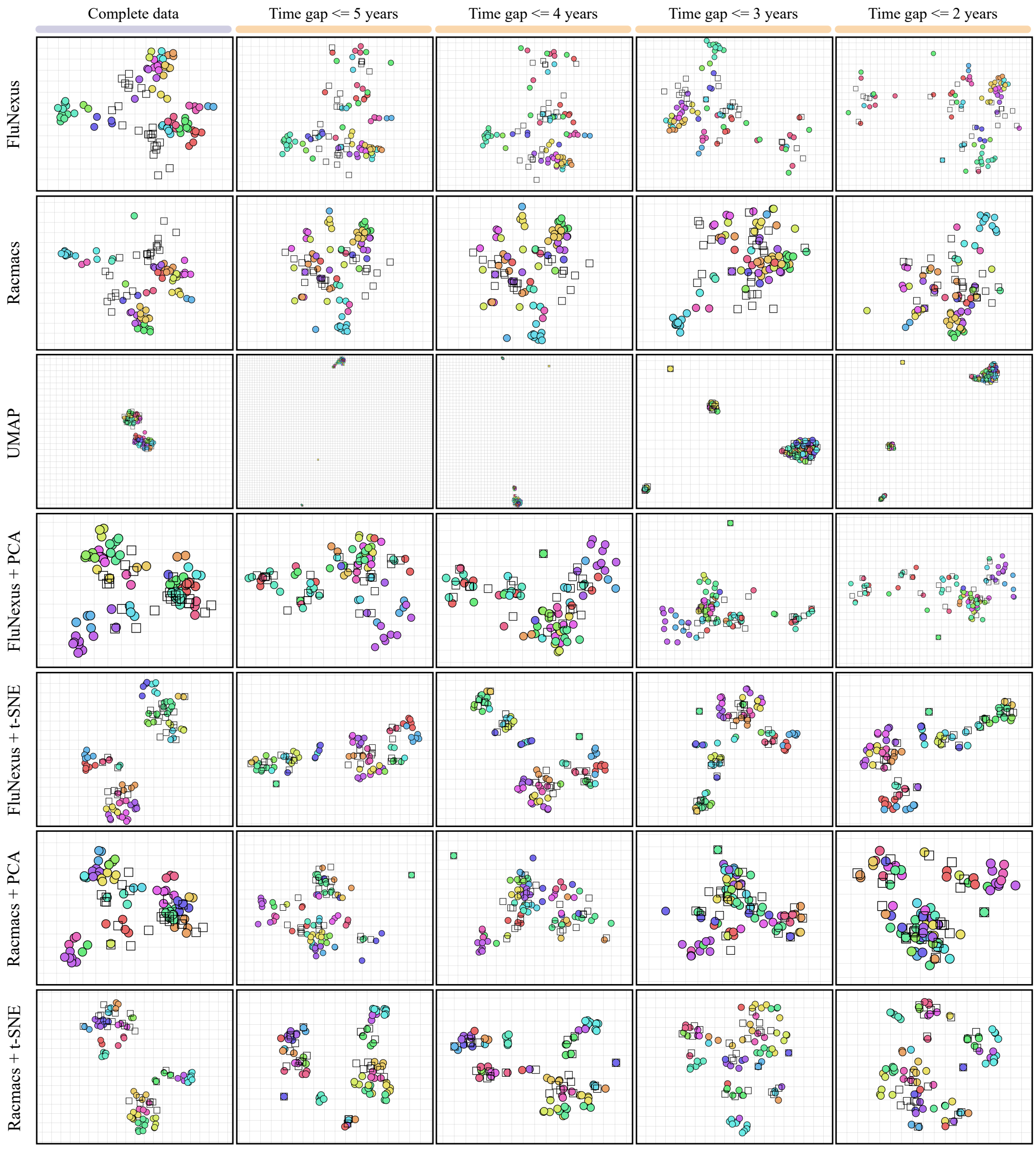


**Figure S6.** Antigenic maps generated by FluNexus, Racmacs, UMAP, PCA, and t-SNE using the HI data of H5 subtype under temporal sampling. For each method, antigens are colored consistent with antigenic map generated from their complete HI data.
